# A unique class of Zn^2+^-binding PBPs underlies cephalosporin resistance and sporogenesis of *Clostridioides difficile*

**DOI:** 10.1101/2022.01.04.474981

**Authors:** Michael D. Sacco, Shaohui Wang, Swamy R. Adapa, Xiujun Zhang, Maura V. Gongora, Jean R. Gatdula, Eric M. Lewandowski, Lauren R. Hammond, Julia A. Townsend, Michael T. Marty, Jun Wang, Prahathees J. Eswara, Rays H.Y. Jiang, Xingmin Sun, Yu Chen

## Abstract

β-Lactam antibiotics, particularly cephalosporins, are major risk factors for *C. difficile* infection (CDI), the most common hospital acquired infection. These broad-spectrum antibiotics irreversibly inhibit penicillin-binding proteins (PBPs), essential enzymes that assemble the bacterial cell wall. Little is known about the *C. difficile* PBPs, yet they play central roles in the growth, infection, and transmission of this pathogen. In this study we discover that PBP2, essential for vegetative growth, is the primary bactericidal target for β-lactams in *C. difficile*. We further demonstrate PBP2 is insensitive to cephalosporin inhibition, revealing a key cause of the well-documented, but poorly understood, cephalosporin resistance in *C. difficile*. For the first time, we determine the crystal structures of *C. difficile* PBP2, which bears several highly unique features, including significant ligand-induced conformational changes and an active site Zn^2+^-binding motif that influences β-lactam binding and protein stability. Remarkably, this motif is shared in two other *C. difficile* PBPs essential for sporulation, PBP3 and SpoVD. While these PBPs are present in a wide range of bacterial taxa, including species in extreme environments and the human gut, they are mostly found in anaerobes, typically Firmicutes. The widespread presence of this convergently evolved thiol-containing motif and its cognate Zn^2+^ suggests it may function as a redox-sensor to regulate cell wall synthesis for survival in adverse environments. Collectively, our findings address important etiological questions surrounding *C. difficile*, characterize new elements of PBP structure and function, and lay the groundwork for antibiotic development targeting both *C. difficile* growth and sporulation.

## Main

*Clostridioides difficile* infection (CDI) is a condition where the anaerobic, spore-forming, bacterium *C. difficile* infects the large intestine, producing symptoms that range from mild-diarrhea to life-threatening colitis. As the most common hospital-acquired infection, the pathogenesis of CDI is well-understood^1^. Initially, antibiotics are administered for an unrelated infection or prophylaxis, causing the gut flora diversity to diminish. Without competition in the large intestine, *C. difficile* can easily proliferate, secreting toxins that cause cell death. The primary risk factor for CDI is the use of broad-spectrum antibiotics, specifically those with weak activity against *C. difficile* and strong activity against other gut bacteria^2^. Currently, 3^rd^ generation cephalosporins and clindamycin present the highest odds ratio for CDI (3.20, 2.86), followed by 2^nd^ and 4^th^ generation cephalosporins (2.23, 2.14)^3^. Other risk factors include treatment duration, age, overall health, and diet. Specifically, recent studies have found Zn^2+^-rich diets increase the likelihood and severity of CDI^4–6^.

Cephalosporins are members of the β-lactam family of antibiotics. These drugs achieve their bactericidal potency by irreversibly inhibiting penicillin-binding proteins (PBPs), particularly the transpeptidase (TPase) PBPs that cross-link cell wall peptidoglycan. Collectively, their enzymatic activity is essential and deactivating individual TPase PBPs can prove lethal to the organism^7,8^. Four annotated TPase PBPs exist in the genome of *C. difficile* R20291, a hypervirulent epidemic RT027 strain: PBP1 (*CDR20291_0712*), PBP2 (*CDR20291_0985*), PBP3 (*CDR20291_1067*), and SpoVD (*CDR20291_2544*)^9^. A recent study found PBP1 and PBP2 are essential for vegetative growth, while SpoVD and PBP3 are essential for sporogenesis. Of these, PBP1 is the only bifunctional (class A) PBP that bears an additional glycosyltransferase (TGase) domain that independently polymerizes the cell wall glycan chain^10^. In contrast, monofunctional TPases (class B PBPs) such as PBP2, PBP3, and SpoVD, must associate with a separate TGase, like those from the SEDS (shape elongation division sporulation) family, to polymerize the glycan units of peptidoglycan^11^.

Cephalosporin resistance in *C. difficile* is well documented, but the underlying mechanism has, until this point, remained unclear^12^. Herein, we use a combination of experimental techniques to characterize the molecular basis of cephalosporin resistance in *C. difficile*. In doing so, we identify a unique role for Zn^2+^ in three *C. difficile* PBPs, which represent a new group of PBPs that have not been previously characterized.

### The essential, monofunctional, PBP2 is the main bactericidal target for β-lactam antibiotics and is poorly inhibited by cephalosporins

By comparing the antibacterial MIC of select members of each β-lactam class (penicillin, cephalosporin, carbapenem, and monobactam) we found resistance is more pronounced amongst cephalosporins (MICs > 64 μg/mL), consistent with previous reports ^13^. Subsequent biochemical analysis of purified PBP1, PBP2, and PBP3 (**Fig. 1a**) demonstrate PBP1 and PBP2 are selectively insensitive to most cephalosporins, in contrast to the non-essential PBP3, which is potently inhibited by all β-lactams. There was a strong correlation between PBP2 inhibition and MIC (r^2^ = 0.735), suggesting PBP2 is the main target through which β-lactams achieve their bactericidal properties against *C. difficile*. (**Fig. 1b**). This correlation is also evident between the MIC and PBP2 IC_50_ values for β-lactams that underwent further analysis (**Fig. 1c, Extended Data Fig. 1**).

**Figure 1.**
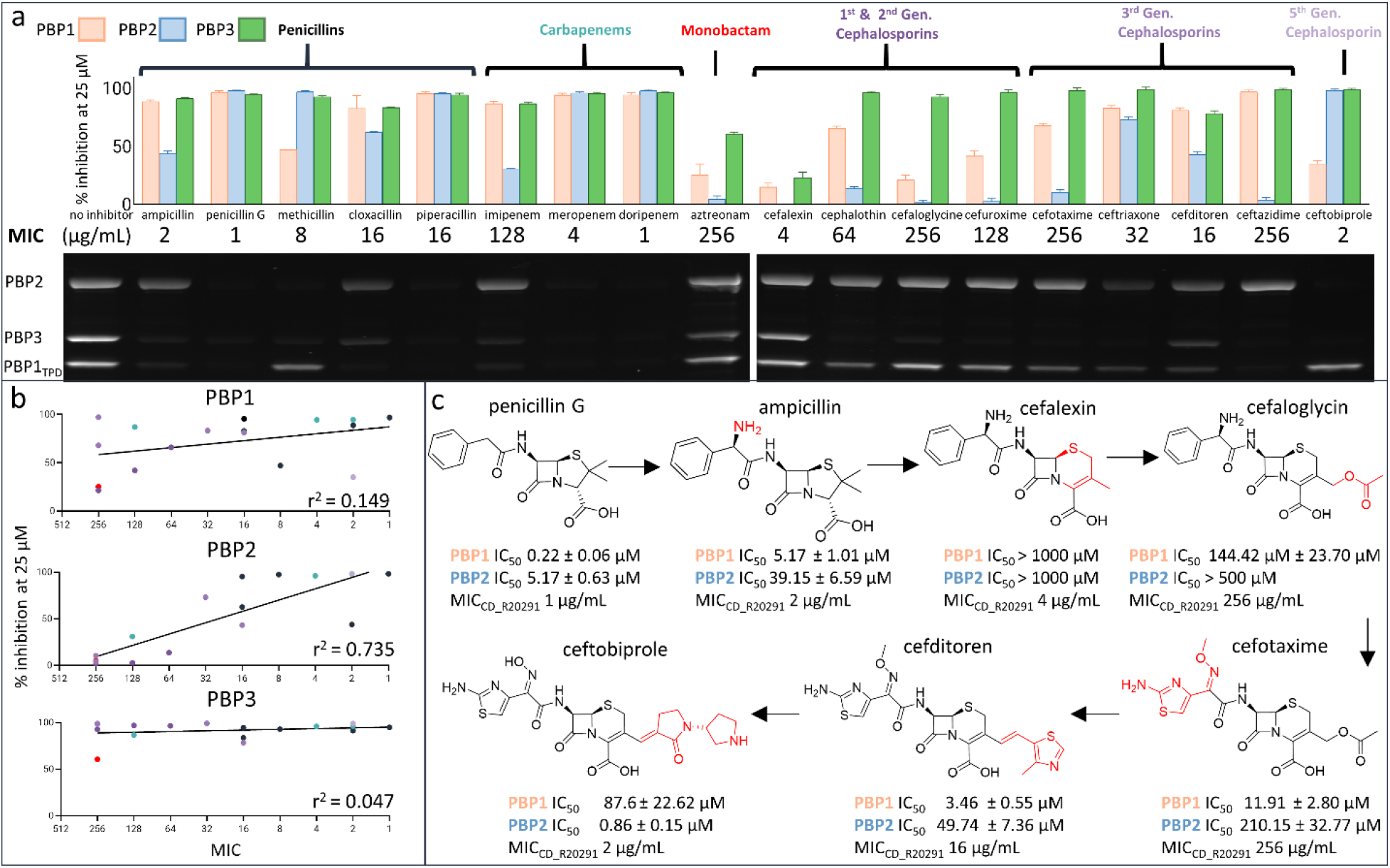
Biochemical and cellular inhibition profile of β-lactam antibiotics against major vegetative *C. difficile* TPase PBPs. **(a)** Single concentration (25 μM) inhibition of fluorescent bocillin binding for select β-lactams against PBP1, PBP2, and PBP3. PBP1 and PBP2 demonstrate cephalosporin insensitivity, which is visible where bands have similar intensity to control (bottom, electrophoresis results, lane 1). **(b)** Relationship between antibacterial potency (MIC) and single concentration % inhibition for each PBP, labelled with its log regression coefficient. Cefalexin was omitted as an outlier. The high coefficient for PBP2 suggests it is the primary bactericidal target of β-lactams. **(c)** IC_50_ values of select β-lactams against PBP1 and PBP2 reveal the cephem nucleus has intrinsically weaker reactivity, but modifications to the sidechain and 3’ leaving group restore potency.

While PBP1 is encoded by an essential gene, there was weak correlation between β-lactam inhibition of PBP1 and antibacterial activity (r^2^ = 0.149), just marginally higher than that of the non-essential PBP3 (r^2^ =0.047). However, because PBP1 is a bifunctional PBP, gene-deletions fail to distinguish whether its TPase or TGase activity is essential. This experiment suggests the TGase domain, and not the TPase domain, probably represents the essential component of this enzyme. Notably, ceftazidime is a selective and potent inhibitor of PBP1, but is non-bactericidal (MIC = 256 μg/mL). Conversely, ceftobiprole was the most potent cephalosporin tested (MIC = 2 μg/mL) and is a selective and potent inhibitor of PBP2 with ~100% inhibition at 25 μM. The inhibition profile of these two cephalosporins independently support the hypothesis that PBP2 is the primary target for β-lactams, while the PBP1 TPase domain is likely non-essential.

### The cephem nucleus of cephalosporins is intrinsically less reactive against PBP1 and PBP2

With few exceptions, cephalosporins poorly inhibit both PBP1 and PBP2, leading us to hypothesize that the cephem nucleus might be intrinsically unreactive towards these enzymes. We subsequently determined the inhibition constants of seven β-lactam drugs, which would answer two outstanding questions, 1) is the cephem nucleus intrinsically less reactive than the penam? 2) If so, can chemical modifications to the 3’ leaving group and sidechain overcome intrinsic barriers to reactivity? A comparison of ampicillin with cefalexin, which is chemically identical except for the cephem nucleus, suggests the cephem nucleus is intrinsically unreactive towards PBP1 and PBP2 (**Fig. 1c**). Cefalexin, however, is atypical since it lacks a 3’ leaving group, which normally contributes to the energetics of cephalosporin binding. By evaluating cephalosporins with different 3’ leaving groups (e.g., cefalogylcin) and side chains (e.g., cefotaxime), we demonstrate that modifications of these functional groups can improve inhibition against PBP1 and PBP2, and consequently, antibacterial potency (**Fig. 1c**). Among them, ceftobiprole was the best inhibitor of PBP2 (IC_50_ = 0.86 ± 0.15 μM) and has unique activity against Gram-positive bacteria expressing low-affinity PBPs, such as MRSA, *E. faecalis, and S. pneumoniae*^14^. Notably, the 3’ vinyl group of ceftobiprole is a poor leaving group and remains adjoined after acylation. Strangely, cefalexin (MIC = 4 μg/mL) has similar antibacterial potency to ceftobiprole (MIC = 2 μg/mL) despite failing to inhibit any of the six TPases tested, including Ldt2 and Ldt3, two enzymes from the cysteine L,D transpeptidase family that also crosslinks cell wall peptides **(Extended Data Fig. 2)**, suggesting it may achieve bactericidal action by cumulative inhibition of non-essential PBPs, or another target altogether.

### Crystal structure of *C. difficile* PBP2 reveals a unique active site zinc-binding motif that contributes to protein stability and activity

*C. difficile* PBP2 is one of the largest known PBPs, with a MW of 111.14 kDa. As such, the primary sequence of PBP2 bears several contiguous regions with no known homology. For the first time, we present its crystal structure with ampicillin at 3.0 Å resolution (**Fig. 2**). Besides being the first published structure of a *C. difficile* PBP, it is the largest PBP published to date (855 residues modelled: 91.37 kDa). The TPase domain adopts a canonical penicilloyl serine transferase fold, characterized by a five stranded β-sheet that is sandwiched by three α-helices. The active site is found between the last β-strand (β3) and five, offset parallel α-helices, forming a narrow substrate-binding cleft, shown here bound to ampicillin through its catalytic Ser492. The second modelled region was the N-terminal domain (NTD, res. 1-110, 241-383), or “pedestal domain”, which has weak sequence, but moderate structural homology to other TPase PBPs^8,15^. The architecture of the NTD is characterized by a“head” of three helices and an “anchor” formed by three β-sheets.

**Figure 2.**
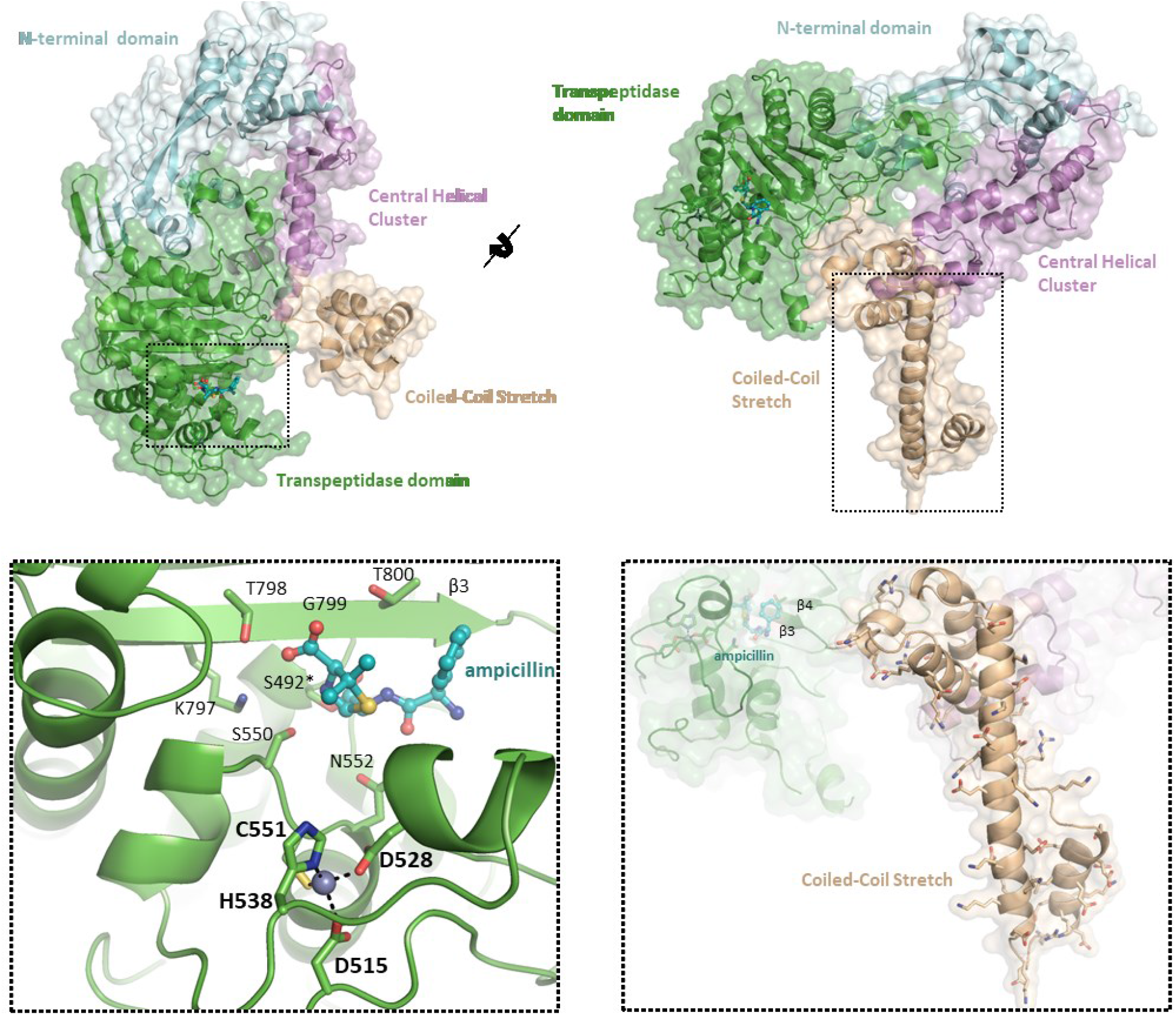
Crystal structure of the PBP2-ampicillin complex. The crystal structure of PBP2 in complex with ampicillin at 3.0 Å resolution reveals three novel structural features. This includes a novel Zn^2+^-binding motif in the active site (bottom left), a charged, elongated, helix connecting the β3-β4 sheets of the TPase domain, termed the “the coiled-coil stretch” (bottom right), and five parallel α-helices, “the central helical cluster”.

PBP2 has two, long stretches of primary sequence with no known homology. The crystal structure shows these regions adopt two well-defined and distinct super-secondary structures that we have termed the “coiled-coil stretch” and the “central helical cluster”. The coiled-coil stretch (res. 805-915), is an elongated coiled-coil that lies orthogonally to the TPase domain. This domain is highly unique because it is found between the β3-β4 sheets of the TPase domain, differing from other published PBPs which usually have an unstructured loop of ~5-20 residues at this position^8,15^. The linchpin of this assembly is an α-helix of 33 residues (**Fig. 2 - bottom right**). Aliphatic residues are concentrated at helix-helix interfaces while a large number of charged residues are mostly in solvent exposed regions, the majority of which are positioned towards the bottom of this helix, suggesting it may interact with the outer leaflet of the membrane bilayer or a separate polar macromolecule. The “central helical cluster” is an insertion (res. 111-240) of five α-helices and two short β-sheets connected by a β-turn between the 1^st^ and 2^nd^ α-helices of the NTD that lies between the coiled-coil stretch and the NTD. Extending throughout the core of PBP2, the central helical cluster forms contacts with all three subregions of the enzyme.

Perhaps the most notable feature of PBP2 is a highly unique Zn^2+^-binding motif adjoined to the active site (**Fig. 2 – bottom left**). Normally concealed in the primary sequence, PBP2 represents the first known instance of an active site Zn^2+^-binding motif in a serine PBP. The Zn^2+^ ion is coordinated by D515, D528, H538, and C551, where C551 is the X residue of the highly conserved SXN motif. The SXN motif plays an important role in substrate binding, suggesting the Zn^2+^ ion is important for maintaining the structural integrity and supporting the catalytic function of the active site^7,8^.

The discovery of a Zn^2+^ in the active site of PBP2, initially through X-ray crystallography **(Fig. 2, Extended Data Fig. 3a)** and subsequently confirmed with native mass spectrometry (**Extended Data Fig. 3b**), led us to investigate its biochemical role. To this end, Cys551 and Asp515 were mutated to Ser and Asn. These mutations (Cys ➔ Ser and Asp ➔ Asn) were designed to displace the Zn^2+^-ion without disrupting structural integrity or generating artificial interactions within the active site. Here we show both C551S and D515N have reduced affinity for the fluorescent penicillin, bocillin (PBP2_WT_*K*_0.5_= 2.95 ± 0.72 μM, PBP2_C551S_*K*_0.5_ = 13.90 ± 1.84 μM, PBP2_D515N_*K*_0.5_ = 16.72 ± 2.72 μM; **Extended Data Fig. 3c**), suggesting the loss of this Zn^2+^ indirectly influences catalytic activity. Furthermore, while WT PBP2 has a melting temperature (T_m_) of 65.45 ± 0.14 °C, C551S and D515N mutants demonstrate a bi-sigmoidal unfolding pattern with reduced T_m_, suggesting denaturation occurs sequentially and independently in two separate domains of the protein (C551S T_m1_ = 36.84 ± 0.17 °C, T_m2_ = 53.71 ± 0.11 °C; D515N T_m1_ = 34.80 ± 0.30 °C, T_m2_ = 53.43 ± 0.11 °C; **Extended Data Fig. 3d**). The altered melting temperature curves for the C551S and D515N mutants suggests this Zn^2+^-binding motif is critical for the structural stability of PBP2.

### Comparison of apo and complex PBP2 structures reveals ligand-induced conformational changes and narrow active site restricting cephem nucleus access

To delineate the structural basis for the cephalosporin insensitivity of PBP2, we solved structures of PBP2 in its un-bound (apo) form (2.8 Å resolution; **Fig. 3a**) and with ceftobiprole (3.0 Å resolution; **Fig. 3b**). When unbound, the active site of PBP2 is occluded and K797, T798, and G799 of the highly conserved KTG motif are distorted. These distortions allow the catalytic serine to adopt a non-productive pose underneath the β3 strand, where it forms a hydrogen bond with a water molecule, making it inaccessible (**Fig. 3a, Extended Data Fig. 4a, b**). However, the active site opens upon ampicillin binding, causing substrate-recognition residues to translate 1-2 Å outwards and the KTG-bearing β3 sheet to shift downward, rigidify, and displace the water molecule. These movements expose the catalytic serine, freeing it for nucleophilic attack. The described active site rearrangements are mostly retained in the ceftobiprole complex **(Fig. 3b, Extended Data Fig. 4c)**, but they are greater than those seen for ampicillin, such as the position of Y529 and adjacent residues, which is consistent with larger size of the cephem nucleus (perimeter = 9.7 Å vs 8.5 Å in ampicillin; **Fig. 3b**). Consequently, the energetic barrier required for binding and acylation may be higher, and additional, compensatory interactions must be formed to overcome this barrier. Thus, without multiple favorable interactions like those of ceftobiprole, most cephalosporins remain weak inhibitors.

**Figure 3.**
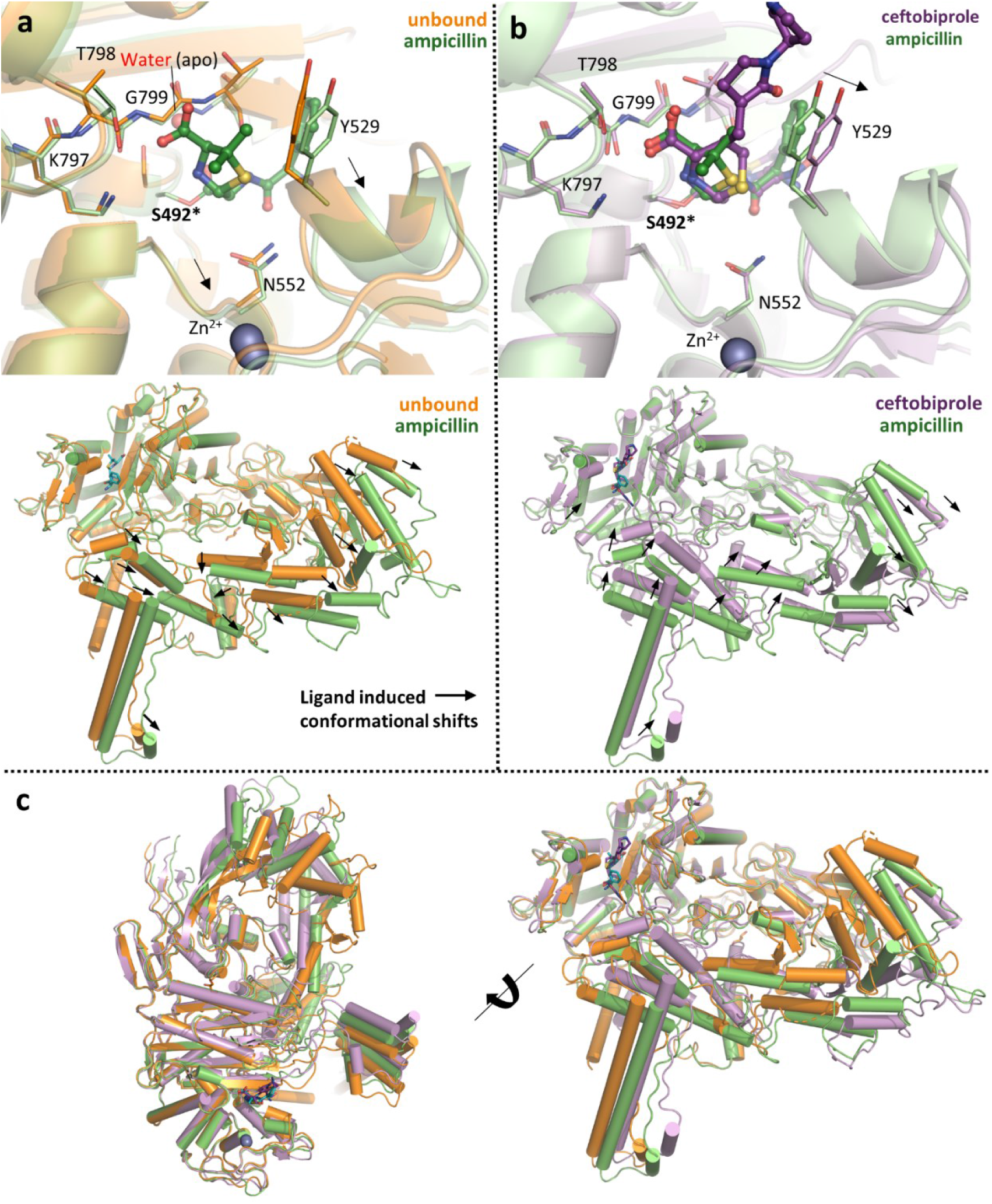
β-lactam binding opens the active site of PBP2 and induces global conformational changes. A superimposition of **(a)** apo-*C. difficile* PBP2 (orange) with ampicillin-bound form (green) shows β-lactam binding causes major conformational changes in the active site (top panel) and peripheral regions (bottom panel). Significant movements are indicated by arrows. **(b)** By comparing the complex structure of ampicillin (green) with ceftobiprole (purple) we find larger conformational changes in the active site (top panel) and overall (bottom panel) that are consistent with the larger size of the cephem core. **(c)** A superimposition of all three structures highlights the global conformations differences in all three structures, with movements mostly localized to the coiled-coil stretch, central-helix cluster, and NTD.

While the active site conformational changes of PBP2 are significant, we observed even larger (> 5-15 Å) rigid body translations outside of the TPase domain (**Fig. 3a, b - bottom panel, Fig. 3c**). These global conformational changes differ between the two complexes and involve regions that likely interact with essential, accessory proteins, including the elongasome protein MreC and a SEDS-family transmembrane TGase MrdB **(Extended Data Fig. 5a, b)**^16,17^. Interestingly, the putative MreC binding cleft formed at the NTD “hinge” splays at different angles between the ampicillin and apo structures, exposing a hydrophobic crevasse that is significantly narrower when bound to ampicillin **(Extended Data Fig. 5c)**. Notably, in the ampicillin bound cleft there is electron density that may correspond to a glycerol that is not present in the apo or ceftobiprole-bound structures. The unique conformations observed for all three structures of PBP2, suggest that occupation of the active site not only induces changes elsewhere in the enzyme, but the identity of the ligand can contribute to different poses. It is possible these movements may promote or negatively impact interactions with partner proteins depending on the nature of the occupying ligand, i.e., substrate (peptidoglycan peptide) or β-lactam. Meanwhile, interactions with protein partners may also influence ligand binding in the active site. Whether and how exactly these conformational changes contribute to regulation of PBP2 activity remains to be investigated. Taken together, the significant local and global conformational changes induced by ligand binding are highly unique for PBP2, even when compared with other β-lactam insensitive PBPs such as *S. aureus* PBP2A^18^, providing the basis for activity regulation and intrinsic resistance to certain β-lactams.

### *C. difficile* and other anaerobic Firmicutes express a novel family of PBPs with an active site Zn^2+^ ion

While certain metallo-β-lactamases and carboxypeptidases rely on Zn^2+^ for catalysis, an active site Zn^2+^ has never been found in a serine TPase PBP^19,20^. Remarkably, in addition to PBP2, this Zn^2+^-binding motif is also present in *C. difficile* PBP3 and SpoVD from *C. difficile*, as well as SpoVD from several other model Firmicutes such as *B. subtilis* (**Fig. 4a**). The presence of Zn^2+^ in *C. difficile* PBP3 and SpoVD, as well as *B. subtilis* SpoVD, was confirmed by native mass spectrometry (**Extended Data Fig. 6)**, meaning three of the four TPase PBPs and all three class B PBPs in *C. difficile* have this motif. Like PBP2, the four residues in the Zn^2+^-binding motif of *C. difficile* PBP3 and SpoVD consist of an invariant His, a Cys from the X residue of the SXN motif, and two electron rich residues -- in this case Cys instead of Asp. They also closely resemble the coordinating residues of the classical Cys_2_His_2_ zinc finger that binds to Zn^2+^ with high affinity^21^. The tight binding of Zn^2+^ in the *C. difficile* PBPs is supported by the observation that PBP2 activity was not affected by 2 mM EDTA and suggests that its presence in these proteins is unlikely an artifact from protein expression and purification in *E. coli*.

**Figure 4.**
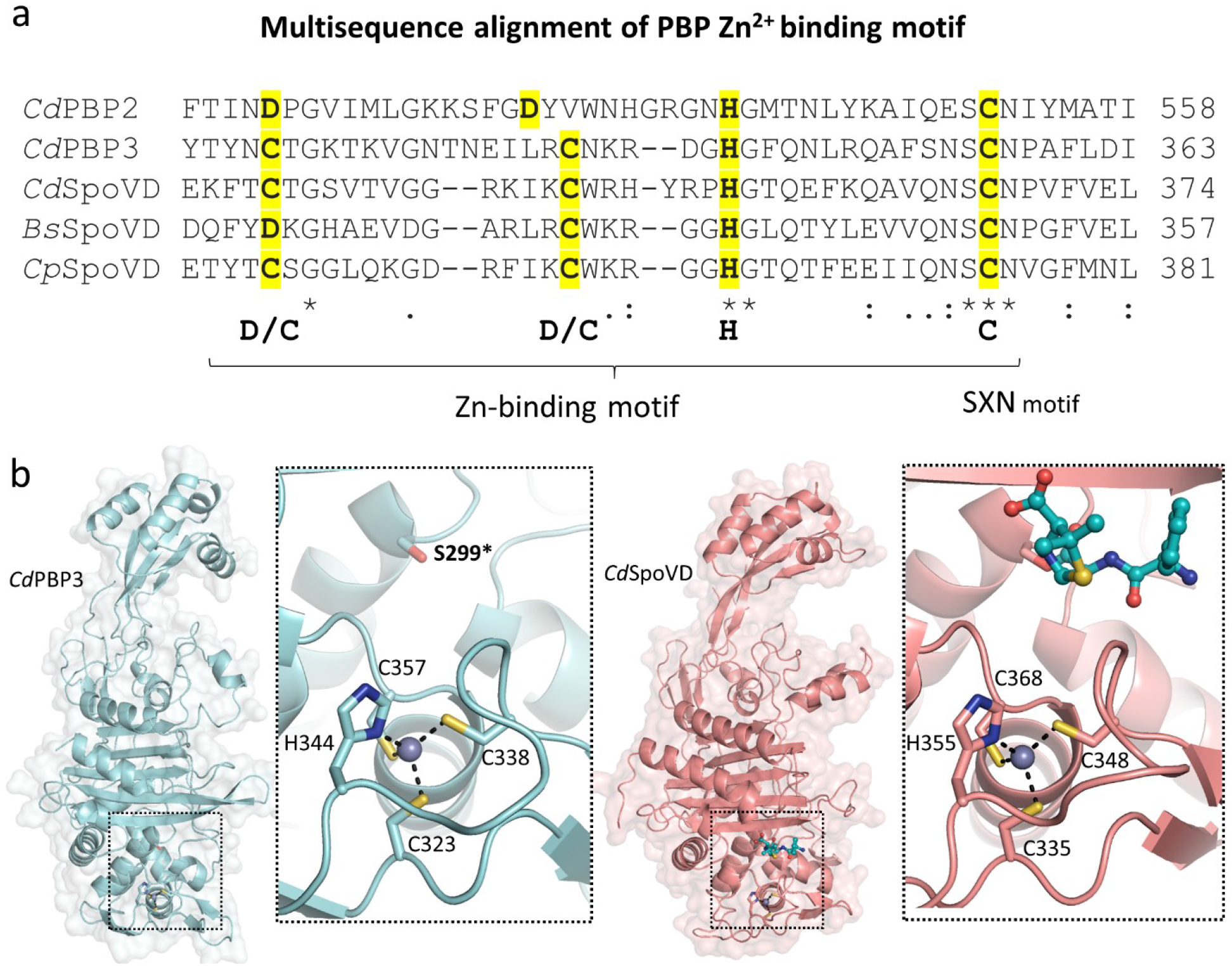
*C. difficile* and other spore-forming Firmicutes express a unique family of Zn^2+^-binding PBPs. **(a)** Multiple sequence alignment uncovers a conserved Zn^2+^ -coordinating tetrad in *C. difficile* PBP2, PBP3, SpoVD and other spore-forming Firmicutes like *B. subtilis* and *C. perfringens* SpoVD. **(b)** The crystal structure of *C. difficile* PBP3 (cyan) and SpoVD (salmon) at 2.40 Å and 2.20 Å reveals a shared fold and Zn^2+^-binding motif.

Next, we determined the structures of PBP3 and SpoVD, at 2.4 Å and 2.2 Å resolution (**Fig. 4b**). Both SpoVD and PBP3 possess a canonical class B PBP fold that closely resembles *P. aeruginosa*and *E. coli* PBP3. Importantly, we find the Zn^2+^ binds in an almost identical manner as PBP2 **(Fig. 2)**, suggesting Asp and Cys are interchangeable for coordinating this ion. The significance of the Asp ➔ Cys substitution is unclear. Whereas the redox-sensitive cysteine in the SXN motif for PBP2 can potentially function as a sensor for environmental oxygen levels, the replacement of Asp for Cys in both sporulation specific PBPs, PBP3 and SpoVD, may further enhance the sensitivity and/or subject them to additional regulation. Previous studies in *B. subtilis* SpoVD have found this motif, which was predicted to be a disulfide bond, interacts with the *B. subtilis* oxidoreductase StoA, suggesting it may function as a unique regulatory mechanism for PBP activity^22^.

### Prevalence of Zn^2+^-binding PBPs in anaerobes through convergent evolution suggests potential adaptation to counter oxygen toxicity

To investigate the presence of Zn^2+^-binding PBPs in a wider range of bacterial species, a hidden markov model (HMM) based on the immediate flanking sequences of the Zn^2+^-binding motif was used to examine the UniProt database (www.uniprot.org). A total of over 5,000 PBPs were retrieved containing the Zn^2+^-binding motif, and a control set of over 5,000 PBPs without the Zn^2+^-binding motif was used for taxa distribution analysis **(Extended Data Table 2)**. The vast majority of bacteria bearing this motif belong to the Firmicutes phylum and most are anaerobes **(Fig. 5a)**. Representative species include those from extreme environments such as deep-sea vents (*Caminicella sporogenes*), oil fields (*Hungateiclostridium thermocellum*), volcanic ashes and hot springs (*Heliobacterium modesticaldum*)^23–25^, suggesting Zn^2+^-binding PBPs might confer survival advantages in adverse conditions, especially anoxic ones.

**Figure 5.**
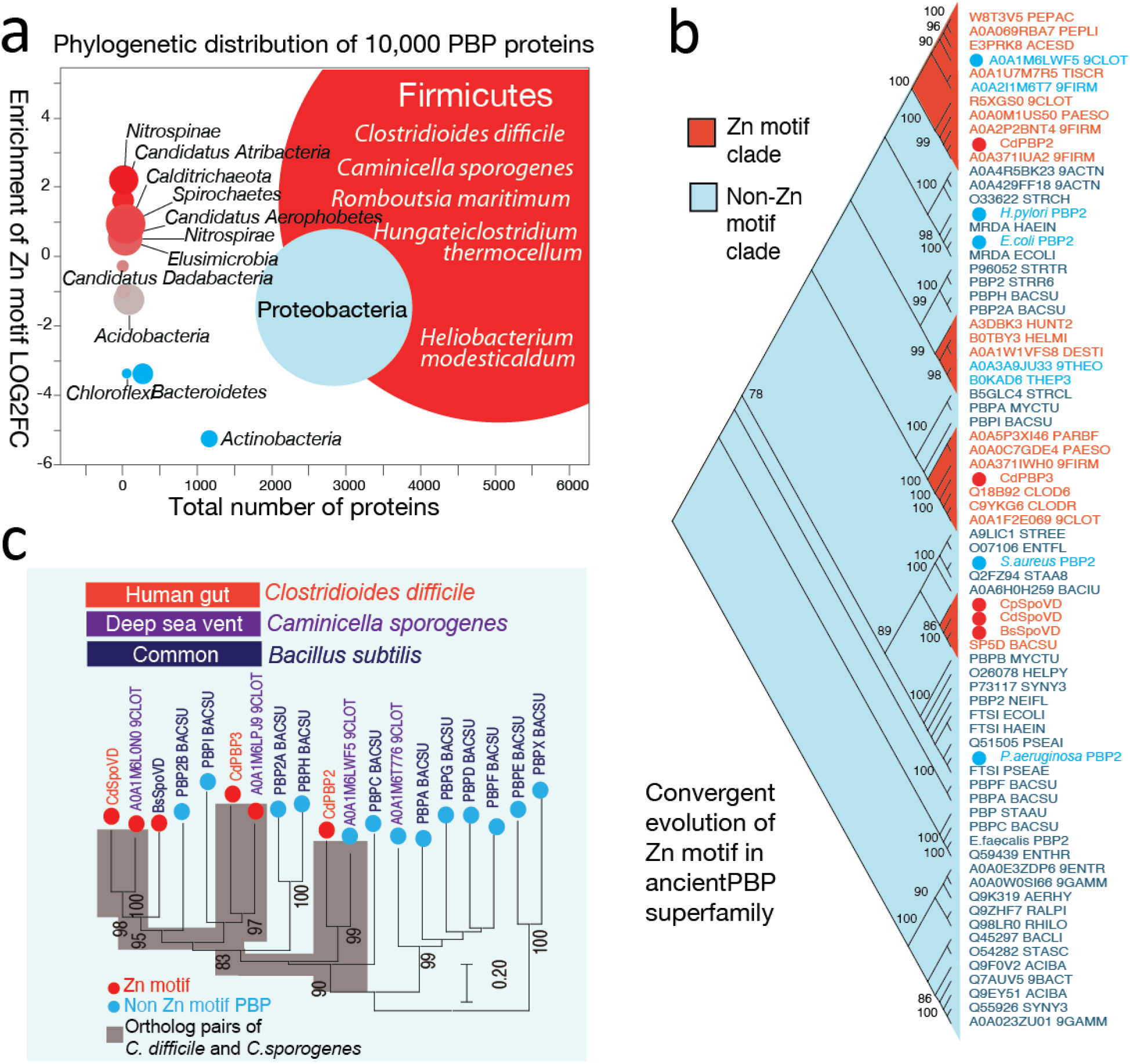
Phylogenetic distributions and evolutionary patterns of Zn^2+-^ binding motifs in the PBP superfamily. Red indicates enrichment of Zn^2+^-binding PBP, blue indicates absence of Zn^2+^-binding PBPs. **(a)** Over 10,000 PBPs were analyzed and Zn^2+-^ binding motifs were identified with a custom-built Hidden Markov Model (HMM). Circle size represents the number of species. Representative species are listed for Firmicutes. **(b)** Structurally diverse PBP superfamily members from diverse taxa were used to build a maximum likelihood tree, revealing Zn^2+^-binding PBPs emerged through convergent evolution in the ancient PBP superfamily. These PBPs have evolved multiple times from independent clades. Bootstrap values are listed next to the branches. **(c)***C. difficile* and the deep-sea vent Gram-negative bacterium *C. sporogenes* share the same set of PBP proteins, two of which (SpoVD and PBP3 homologs) bear Zn^2+^-binding motifs.

Next, a set of phylogenetically diverse PBPs based on the PFAM seed PBP domain collections (Transpeptidase; PF00905) were selected to examine the evolutionary pattern of Zn^2+-^ binding PBPs **(Extended Data Table 3)**. Unexpectedly, we find the Zn^2+^-binding motif independently evolved through convergent evolution from the ancient PBP superfamily, as shown by the four clades in the phylogenetic tree **(Fig. 5b)**. Interestingly, in the PBP2 clade, there are two proteins without a Zn^2+^-binding motif, suggesting it can be gained or lost in a relatively short evolutionary span. For example, *C. sporogenes*, an anaerobic, Gram-negative bacterium isolated from a deep-sea vent shares a similar set of PBPs (PBP2, PBP3 and SpoVD) with *C. difficile* **(Fig. 5c)**. Although *C. sporogenes* PBP3 and SpoVD possess a Zn^2+^-binding motif, its PBP2 orthologue (Uni-prot A0A1M6LWF5_9CLOT) does not, despite its high sequence (60.3%) and predicted structural similarity to *C. difficile* PBP2 **(Extended Data Fig. 7)**. This degree of homology suggests *C. sporogenes* PBP2 can acquire the capacity for Zn^2+^-binding capacity through one or two mutations. Furthermore, because *C. difficile* PBP2 is implicated in vegetative growth and PBP3/SpoVD facilitate sporulation, the Zn^2+^-binding motif may be required for different biological processes, depending on the specific bacterium and the natural habitat.

Notably, bacteria containing Zn^2+^-binding PBPs within the PBP2 clade are exclusively anaerobic. Considering that cysteine residues, including those involved in Zn^2+^-binding, are involved in redox sensing in other proteins ^26,27^, we hypothesize that the Zn^2+^-binding PBPs may function as a regulatory mechanism that can shut down cell wall synthesis or promote sporulation when confronted with molecular oxygen and/or reactive oxygen species, in conjunction with the previously studied transcription-level responses^28^. This hypothesis is partly supported by the afore-mentioned observation that *B. subtilis* SpoVD is regulated by the oxidoreductase StoA^*22*^. Furthermore, as many antibiotics achieve at least part of their bactericidal effects through reactive oxygen species ^29^, such mechanisms may possibly contribute to antibiotic resistance in bacteria such as *C. difficile*.

### All classes of β-lactam antibiotics and the Zn^2+^-chelators calprotectin and TPEN inhibit *C. difficile* sporulation

Several lines of evidence suggest dietary Zn^2+^ increases the severity and likelihood of CDI, but the pleiotropic roles of Zn^2+^ have obfuscated the exact mechanism by which it facilitates pathogenesis. Recent studies have found that calprotectin, a protein secreted by the immune system in response to intestinal inflammation, inhibits *C. difficile* growth^5^. Interestingly, calprotectin is a chelator of divalent cations such as Ca^2+^ and Zn^2+^. Using calprotectin and the hexadentate ligand TPEN we show sporulation can also be significantly inhibited at 10 μg/mL and 12.5 μM (**Fig. 6a**). This anti-sporulation effect is reversed by supplementing the media with Zn^2+^, suggesting this effect is specific to Zn^2+^, even though most certainly not limited to PBP activity. The consistent presence of a Zn^2+^ motif in all three class B *C. difficile* PBPs – PBP2, PBP3, and SpoVD, and its apparent importance for enzymatic activity and sporulation, suggests Zn^2+^ may be uniquely associated with growth of the vegetative cell and spore-cortex. Furthermore, these PBPs may have evolved to take advantage of the anaerobic environment of the colon, which is rich in excreted trace metals such as Zn^2+ 30^. Thus, while Zn^2+^ likely plays multiple important roles in *C. difficile* growth, it at least impacts sporulation and may contribute to the relationship between CDI and colonic Zn^2+^ concentration.

**Figure 6.**
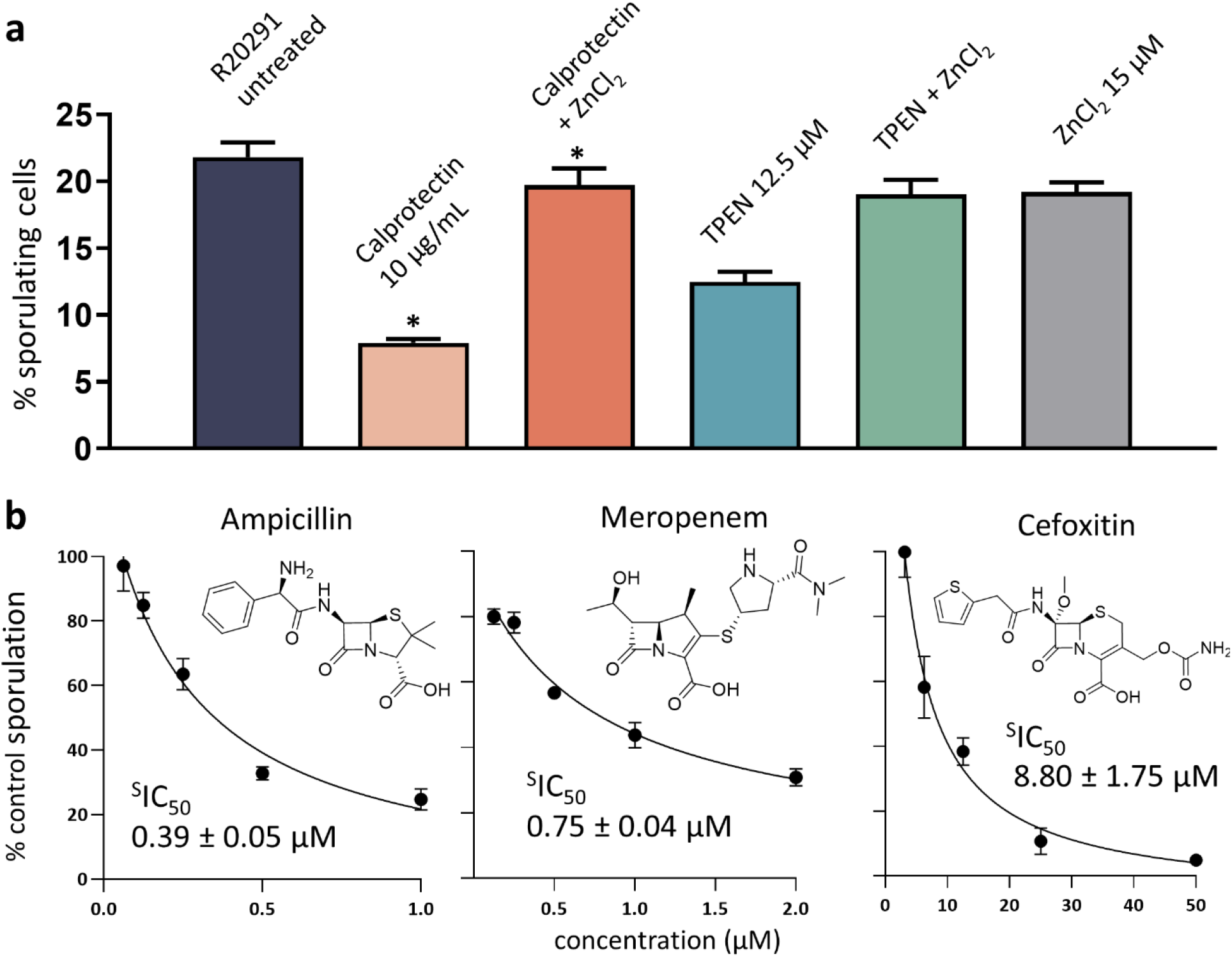
Zn^2+^-chelating agents TPEN, calprotectin and all classes of β-lactams disrupt *C. difficile* sporulation. **(a)** Anti-sporulation effects of the chemical chelator TPEN and the biological chelator calprotectin, an innate immune system protein. This inhibition can be reversed through restoring growth media with Zn^2+^ (RM one-way ANOVA, * P < 0.05, nonsignificant where undesignated). **(b)** All three major classes of β-lactams-ampicillin (penicillins), meropenem (carbapenems), and cefoxitin (cephalosporins)- were assessed at sub-MIC concentrations (½ MIC), revealing these antibiotics are potent inhibitors of sporulation (indicated by sporulation IC_50_, or ^S^IC_50_) via inhibition of PBPs required for spore cortex assembly, such as PBP3 and SpoVD.

Other anti-sporulating agents include cephamycins, a class of cephalosporins that inhibit the enzymatic activity of SpoVD. Because PBPs are integral for spore-cortex formation, we hypothesized that, in addition to cephamycins, all β-lactam antibiotics should also reduce sporulation. Here we show that ampicillin (sporulation IC_50_, or ^s^IC_50_, 0.39 ± 0.05 μM) and meropenem (^s^IC_50_ 0.75 ± 0.04 μM) are more efficacious than cefoxitin (^s^IC_50_ 8.80 ± 1.75 μM) *in vitro* (**Fig. 6b**). Although the use of β-lactams to inhibit *C. difficile* sporulation is not practical, they may be uniquely effective against other pathogens such as *C. perfringens* and *B. anthracis*. Nonetheless, SpoVD and PBP3 are very promising drug targets. A selective, non-bactericidal inhibitor of these enzymes could be paired with the standard treatment regimen, thus mitigating the likelihood of recurrent infection and transmission associated with spore formation.

## Conclusion

Independently, PBPs and CDI are heavily studied subjects, yet very little research exists for the PBPs of *C. difficile*. This study helps delineate the molecular basis for cephalosporin resistance in *C. difficile* and the high risk-factors associated with these antibiotics. More specifically, we demonstrate that *C. difficile* PBP1 and PBP2 are poorly inhibited by most cephalosporins, resulting in lower antibacterial activity. The insensitivity of *C. difficile* to cephalosporins, coupled with broad-spectrum activity against gut bacteria, contribute to the extraordinary high odds-ratio associated with this class of antibiotics. Our discovery of a new group of Zn^2+^-binding PBPs offers further insights into the Zn^2+^-dependence of *C. difficile*, while identifying a potential regulatory module for cell wall synthesis that is present in a wide range of bacteria. Collectively, the characterization of these PBPs leads to new and important questions about the role and regulation of these PBPs, their novel domains, and the Zn^2+^-binding motif as it relates to vegetative cell growth, division, and sporogenesis in this unique pathogen. The evaluation of their inhibition profiles, relative essentiality, and crystal structures also provides a structural framework for future antibiotic discovery against *C. difficile*, especially considering the recent headway of novel non-β-lactam chemical scaffolds as PBP inhibitors ^31^.

## Materials and Methods

### Protein cloning and purification

The coding sequence corresponding to the TPase domain of PBP1 (*CDR20291_0712*), the soluble constructs of PBP3 (*CDR20291_1067*), Ldt2 (*CDR20291_2601*), Ldt3 (*CDR20291_2843*), PBP2 (*CDR20291_0985*), SpoVD *CDR20291_2544*, and the crystallization constructs of PBP2 (*CDR20291_0985*) and PBP3 (*CDR20291_1067*), were amplified from *C. difficile* R20291 genomic DNA^9^. The soluble sequence of BSU_15170 (BsSpoVD) was amplified from the genomic DNA of *B. subtilis* strain 168^32^ The expression vector was modified from pETGST, in which the thrombin cleavage site was replaced by the TEV (named pETGSTTEV) or ULP1 (named pETGSTSUMO) cleavage site. The vector was digested and the PCR fragments for each protein was inserted into the multi-clone site accordingly. PBP1 376-765 was inserted into NheI/HindIII site; PBP2 36-962 and Ldt2 full length were inserted into NheI/XhoI sites; Ldt3 was inserted into the NdeI/XhoI site. SpovD 38-583 was inserted into the NheI/XhoI site of the pETGSTSUMO vector. PBP2 D515N and C551S mutants were generated using the QuikChange Lightning Site-Directed Mutagenesis Kit (Agilent Technologies). Cloned vector was then transformed into BL21(DE3) pLysS *E. coli*. Standard overnight cultures were grown in LB media containing chloramphenicol and kanamycin and used to inoculate 1 L LB cultures. Cultures were grown until the optical density (OD_600_) reached ~0.7. Protein expression was then induced with 0.5 mM isopropyl-β-d-thio-galactopyranoside (IPTG) overnight at 20 °C. Cells were harvested via centrifugation at 4000 x g for 10 minutes at 4 °C. The cell pellet was suspended with a solution of 20 mM Tris pH 8.0, 200 mM NaCl, 20 mM imidazole, and one dissolved Thermo Scientific^™^ Pierce Protease Inhibitor Tablet. Cells were lysed with a sonicator using a 10 second sonication/15 second rest cycle for 15 minutes. Cell lysate was centrifuged at 45,000 x g for 35 minutes and supernatant was loaded onto a HisTrap HP affinity column (GE Healthcare). A linear gradient of buffer B (20 mM Tris pH 8.0, 300 mM NaCl, 500 mM imidazole, and 10% glycerol) was applied to elute the recombinant protein, which usually occurred at 30% buffer B. These fractions were pooled, and buffer exchanged three times into protease buffer (20 mM Tris pH 8.0, 10% glycerol) using an Amicon Ultra centrifugal filter (Sigma-Aldrich). Purified His6-tagged TEV protease was added to all proteins at a 1:20 ratio (except for SpoVD) and incubated overnight at 4 °C. Pure His6-tagged ULP1 protease was added to SpoVD at 1:20 ratio and incubated overnight at 4 °C. The digested protein was re-loaded onto the HisTrap HP affinity column. Flowthrough was collected, concentrated, and loaded to a HiLoad 16/60 Superdex 75 size exclusion column (GE Healthcare) where it ran at a flow rate of 0.5 mL/min. Peak fractions were combined and purity assessed (> 95%) with gel-electrophoresis. The identity of each enzyme was subsequently characterized with gelelectrophoresis, native-mass spectrometry, bocillin or nitrocefin binding, and/or X-ray crystallography.

### X-ray crystallography

8 mg/mL PBP2 was incubated with 1 mM ampicillin for 2 hours at room temperature. Crystals were grown by mixing 1.5 μL of the protein + inhibitor solution with 1.5 μL of well solution: 15% PEG 4000, 0.2 M ammonium sulfate, and 0.1 M sodium citrate pH 5.6. Cover slips containing the drops were sealed and equilibrated over 1 mL of well solution for one week at 20 °C, producing many plate-like crystals. These crystals were then crushed into seeds. This method was repeated, but instead of mixing the well stock with protein, diluted seed stock was used to control the rate of nucleation, yielding diffraction quality crystals. Crystals were then briefly soaked in a cryoprotectant solution of 30% PEG 4000, 0.2 M ammonium sulfate, 0.1 M sodium acetate pH 5.6, 15% glycerol and flash frozen in liquid nitrogen.

Surprisingly, PBP3 readily crystallized in the same conditions as PBP2 (15% PEG 4000, 0.2 M ammonium sulfate, 0.1 M sodium citrate pH 5.6). However, they were highly sensitive to glycerol in the cryoprotectant solution. To address this, crystals were crushed into seeds and diluted into the crystallization solution supplemented with 5% glycerol. Crystals were regrown by mixing 12 mg/mL protein with the 5% glycerol diluted seed stock. After reaching full size in a week, 2 μL of a cryoprotectant solution containing 25% PEG 4000 and 30% glycerol was added directly to the drops for ~20 seconds and crystals were flash-frozen in liquid nitrogen.

### Model building and refinement

X-ray diffraction data was collected on the Structural Biology Center (SBC) 19-ID and Southeast Regional Collaborative Access Team (SER-CAT) 22-ID beamlines at the Advanced Photon Source in Argonne, IL and processed and scaled with the CCP4 versions of iMosflm and Aimless^33–35^. Initial models were obtained using the MoRDa package of the online CCP4 suite^36^. Models were then processed with PHENIX Autobuild^37,38^. Unsolved regions were manually traced with a polyalanine backbone using COOT and refined with refmac5^39,40^. Initially, aided with the secondary-structure prediction software PSIPRED^41^, α-helices were generated by tracing the electron density map, and bulky, aromatic sidechains such as tryptophan were fit into their corresponding positions. This allowed us to solve the sequence of adjacent residues. Iterations were generated until individual sidechains could be resolved. Two flexible regions with ambiguous electron density were left unmodelled: 357-370 of the NTD and 624-692 of the TPase domain. This region in the TPase domain likely contains three parallel α-helices, but we were unable to confidently model the density. Likewise, a short segment of 197-216 in the NTD was left unmodelled for PBP3. All structural images were generated using PyMol (Schrödinger, LLC).

### Bocillin inhibition assay

Reactions were performed in a 96-well polystyrene plate with a final volume of 60 μL at 37 °C in 1X Tris-buffered saline (20 mM Tris, 150 mM NaCl). For IC_50_ values, inhibitor was serially diluted with a two-fold scheme so the highest concentration had a final value of 1 mM or 500 μM, and the final concentration was either 0.97 μM or 0.489 μM. Single concentration screens were performed at 25 μM. Protein was then diluted and added to the well for a final concentration of 1 μM and incubated for 15 minutes at 37 °C. 1 μL of bocillin (BOCILLIN^™^ FL Penicillin, Sodium Salt, Thermo Fisher Scientific) was added to each well for a final concentration of 20 μM, and incubated for 10 minutes at 37 °C. Reactions were killed with 10 μL of 6X SDS loading buffer and pooled or directly loaded onto a 15-well Novex Wedgewell 8% Tris-Glycine Gel (Thermo Fisher Scientific) and ran at 175 V for ~ 1 hour. Gels were then imaged with a Gel Doc XR+ imager (Bio-Rad Laboratories) for 6 seconds with the fluorescein blot filter setting and analyzed using ImageJ^42^. All values were normalized to background intensity and divided by the average of three control intensities (no inhibitor) for % inhibition value. Assays were performed in duplicate or triplicate and IC_50_ values were calculated using Graphpad Prism with a 3-parameter logistic fit.

### Nitrocefin inhibition assay

Reactions were performed in a 96-well polystyrene plate with a final volume of 60 μL at 37 °C in 20 mM Tris pH 7, 200 mM NaCl). Inhibitors were diluted for a final concentration of 25 μM. Protein was then added to the well for a final concentration of 1 μM and incubated for 15 minutes at 37 °C. 1 μL of nitrocefin (Millipore Sigma) was then added to each well for a final concentration of 20 μM, and reaction progress was monitored using a BioTek Cytation 5 plate reader (Bio-Tek Instruments) at 488 nm and 37°C for 1 hour. Reaction rates were normalized to the control (no inhibitor) for % inhibition values. Assays were performed in duplicate and mean % inhibition values were calculated using Graphpad Prism with a 3-parameter logistic fit.

### Thermal shift assay

Thermal shift assays were performed using differential scanning fluorometry. Purified protein was diluted in TBS and combined with SYPRO^™^ Orange (Invitrogen) for a final protein concentration of 3 μM and final SYPRO^™^ Orange concentration of 1X, with a final volume of 20 μL. Fluorescence was measured using The Applied Biosystems^®^ 7900HT Fast Real-Time PCR (Thermo Fisher Scientific) from 25 °C to 99 °C with a ramp step of 0.3 °C. Melting temperature values were determined by fitting sigmoidal fluorescence intensity curves with a 4-parameter logistic fit.

### Minimum inhibitory concentration (MIC) determination

The MICs of antibiotics on *C. difficile* strain R20291 were determined as described previously^43^. Antibiotics were added to wells of 96-well microplates containing R20291 cultures at a density of 0.5 McFarland (100 μL per well) in BHI medium to make final concentrations of antibiotics ranging from 128 μg/mL to 0.5 μg/mL at a two-fold dilution. The plates were incubated at 37 °C for 24 hours, and MIC was determined as the lowest concentration that completely inhibits the bacterial growth in the wells.

### Sporulation assays

Sporulation assays were performed as described previously with minor modifications^44–46^.

Briefly, *C. difficile* R20291 cells were cultured to mid-log phase in BHIS medium supplemented with 0.1% taurocholate (Sigma) at 37 °C in an anaerobic chamber. A mixture of 70:30 sporulation medium (70% SMC medium and 30% BHIS medium containing 63 g Bacto peptone, 3.5 g protease peptone, 11.1 g BHI medium, 1.5 g yeast extract, 1.06 g Tris base, 0.7 g ammonium sulfate, and 15 g agar per liter) was prepared. Cultures were subsequently diluted to an optical density of 0.5 at OD600, then 150 ul of R20291 cultures were spread over the surface of a 70:30 medium agar plate. Approximately 24 hours after the start of stationary phase (T24), *C. difficile* cells were scraped from the surface of the plate with a sterile inoculating loop and suspended in approximately 5 mL BHIS to an OD_600_ = approximately 1.0. To assess ethanol-resistant spore formation, 500 μL of samples from the sporulation medium were removed from the anaerobic chamber and mixed 1:1 with 95% ethanol for 15 minutes to kill vegetative cells. The samples were then returned to the anaerobic chamber, 100 μl of the ethanol-treated cultures were mixed with 100 μl of 10% taurocholate, and the mixtures were plated onto BHIS agar to induce *C. difficile* spore germination. The ethanol resistant CFU/mL was determined after incubation for 24 hours and was divided by the total CFU/mL of the non-ethanol treated cultures. A minimum of three biological replicates were performed.

### PBP sequence analysis

To examine the common protein domain families for discovery of patterns of site-specific evolutionary conservation related to Zn2+-binding PBPs, profile hidden Markov models (profile HMMs) were built for probabilistic domain searches. HMMER (v.3.3.2) uses ensemble algorithms to consider all possible sequences alignments, each weighted by likelihood of positive scores. The sequences used as an input are listed in Extended Data Table 3. The databases of UniProtKB and SwissProt were used as targets for the search. For phylogenetic analysis, a set of most representative ‘seeds’ sequences were selected from the superfamily of over 200,000 members, the phylogenetically diverse domains of DD-transpeptidase (PF00905) were used analyze the evolutionary patterns.

### Native mass spectrometry

All proteins were buffer exchanged into 0.2 M ammonium acetate using Bio-Spin P-6 Size Exclusion Spin Columns (BioRad) as previously described^47^. Native MS was performed using a Q-Exactive HF quadrupole mass spectrometer with Ultra-High Mass Range research modifications (Thermo Fisher Scientific). All proteins were ionized using nano-electrospray ionization in positive mode using 1.1-1.5 kV of spray voltage at 200 °C. Samples were analyzed with a 2,000-15,000 *m/z* range and the resolution was set to 15,000. The trapping gas was set to 5 for all samples and 50 V was applied in the source to aid in desolvation. Data were deconvolved and analyzed using UniDec^48^.

## Data Availability

All crystal structures have been deposited in the RCSB Protein Data Bank (PDB) with accession IDs of: *C. difficile* PBP2 + ampicillin (PDB ID 7RCW), unbound PBP2 (PDB ID 7RCX), PBP2 + ceftobiprole (PDB ID 7RCY), SpoVD + ampicillin (PDB ID 7RCZ), and PBP3 (PDB ID 7RD0).

## Acknowledgements

We thank Sean Crosson, Phoebe Rice and Gloria Ferreira for their consult. We also thank the staff members of the Advanced Photon Source of Argonne National Laboratory, particularly those at the Structural Biology Center (SBC) and Southeast Regional Collaborative Access Team (SERCAT), for X-ray diffraction data collection. SBC-CAT is operated by UChicago Argonne LLC, for the U.S. Department of Energy, Office of Biological and Environmental Research under contract DE-AC02-06CH11357. The use of beamlines at SER-CAT was supported by its member institutions, and equipment grants (Grants S10_RR25528 and S10_RR028976) from the NIH.

## Author contributions

The studies presented herein were conceived and designed by M.D.S., X.S., and Y.C.; The manuscript was written by M.D.S. and Y.C. with input from others; J.W., M.T.M., P.J.E, R.J., X.S., and Y.C. provided scientific input, funded and supervised the studies; Figures were prepared by M.D.S. with input from others; Protein constructs were designed by M.D.S. and Y.C.; L.R.H. assisted with the experiments on *B. subtilis* SpoVD including its cloning with input from P.J.E. Proteins were cloned and purified by X.Z.; Biochemical and thermal shift binding assays were performed by M.D.S.; Crystallization experiments were performed by M.D.S. with help from M.V.G. and J.R.G.; Crystal structures were solved and refined by M.D.S. with help from E.M.L.; J.A.T. performed native mass spectrometry experiments with guidance from M.T.M.; R.J. and S.R.A. performed phylogenetic analysis and prepared Fig. 5; MIC and sporulation assays were performed by S.W.

## References

1 Leffler, D. A. & Lamont, J. T. Clostridium difficile infection. New England Journal of Medicine 372, 1539–1548 (2015).

2 Bartlett, J. G. Historical perspectives on studies of Clostridium difficile and C. difficile infection. Clinical infectious diseases 46, S4–S11 (2008).

3 Slimings, C. & Riley, T. V. Antibiotics and hospital-acquired Clostridium difficile infection: update of systematic review and meta-analysis. Journal of Antimicrobial Chemotherapy 69, 881–891 (2014).

4 Zackular, J. P. et al. Dietary zinc alters the microbiota and decreases resistance to Clostridium difficile infection. Nature medicine 22, 1330–1334 (2016).

5 Zackular, J. P. & Skaar, E. P. The role of zinc and nutritional immunity in Clostridium difficile infection. Gut microbes 9, 469–476 (2018).

6 Lopez, C. A. et al. The immune protein calprotectin impacts Clostridioides difficile metabolism through zinc limitation. Mbio 10, e02289–19 (2019).

7 Sauvage, E. & Terrak, M. Glycosyltransferases and transpeptidases/penicillin-binding proteins: valuable targets for new antibacterials. Antibiotics 5, 12 (2016).

8 Sauvage, E., Kerff, F., Terrak, M., Ayala, J. A. & Charlier, P. The penicillin-binding proteins: structure and role in peptidoglycan biosynthesis. FEMS microbiology reviews 32, 234–258 (2008).

9 Stabler, R. A. et al. Comparative genome and phenotypic analysis of Clostridium difficile 027 strains provides insight into the evolution of a hypervirulent bacterium. Genome biology 10, 1–15 (2009).

10 Lovering, A. L., de Castro, L. H., Lim, D. & Strynadka, N. C. Structural insight into the transglycosylation step of bacterial cell-wall biosynthesis. Science 315, 1402–1405 (2007).

11 Meeske, A. J. et al. SEDS proteins are a widespread family of bacterial cell wall polymerases. Nature 537, 634–638 (2016).

12 Spigaglia, P. Recent advances in the understanding of antibiotic resistance in Clostridium difficile infection. Therapeutic advances in infectious disease 3, 23–42 (2016).

13 Khanafer, N. et al. Susceptibilities of clinical Clostridium difficile isolates to antimicrobials: a systematic review and meta-analysis of studies since 1970. Clinical Microbiology and Infection 24, 110–117 (2018).

14 Bogdanovich, T., Ednie, L. M., Shapiro, S. & Appelbaum, P. C. Antistaphylococcal activity of ceftobiprole, a new broad-spectrum cephalosporin. Antimicrobial agents and chemotherapy 49, 4210–4219 (2005).

15 Macheboeuf, P., Contreras-Martel, C., Job, V., Dideberg, O. & Dessen, A. Penicillin binding proteins: key players in bacterial cell cycle and drug resistance processes. FEMS microbiology reviews 30, 673–691 (2006).

16 Contreras-Martel, C. et al. Molecular architecture of the PBP2-MreC core bacterial cell wall synthesis complex. Nature communications 8, 776 (2017).

17 Sjodt, M. et al. Structural coordination of polymerization and crosslinking by a SEDS-bPBP peptidoglycan synthase complex. Nature Microbiology 5, 813–820 (2020).

18 Lim, D. & Strynadka, N. C. J. Structural basis for the β lactam resistance of PBP2a from methicillin-resistant Staphylococcus aureus. Nature Structural Biology 9, 870–876 (2002).

19 Dideberg, O. et al. Structure of a Zn 2+-containing D-alanyl-D-alanine-cleaving carboxypeptidase at 2.5 Å resolution. Nature 299, 469–470 (1982).

20 Bush, K. Metallo-β-lactamases: a class apart. Clinical infectious diseases 27, S48–S53 (1998).

21 Laity, J. H., Lee, B. M. & Wright, P. E. Zinc finger proteins: new insights into structural and functional diversity. Current Opinion in Structural Biology 11, 39–46 (2001).

22 Liu, Y., Carlsson Möller, M., Petersen, L., Söderberg, C. A. & Hederstedt, L. Penicillin-binding protein SpoVD disulphide is a target for StoA in Bacillus subtilis forespores. Molecular microbiology 75, 46–60 (2010).

23 Kimble, L. K., Mandelco, L., Woese, C. R. & Madigan, M. T. Heliobacterium modesticaldum, sp. nov., a thermophilic heliobacterium of hot springs and volcanic soils. Archives of Microbiology 163, 259–267 (1995).

24 Alain, K. et al. Caminicella sporogenes gen. nov., a novel thermophilic spore-forming bacterium isolated from an East-Pacific Rise hydrothermal vent. International journal of systematic and evolutionary microbiology 52, 1621–1628 (2002).

25 Zhang, X. et al. Petroclostridium xylanilyticum gen. nov., sp. nov., a xylan-degrading bacterium isolated from an oilfield, and reclassification of clostridial cluster III members into four novel genera in a new Hungateiclostridiaceae fam. nov. International journal of systematic and evolutionary microbiology 68, 3197–3211 (2018).

26 Hübner, C. & Haase, H. Interactions of zinc- and redox-signaling pathways. Redox biology 41, 101916 (2021).

27 Giles, N. M. et al. Metal and redox modulation of cysteine protein function. Chemistry & biology 10, 677–693 (2003).

28 Lu, Z. & Imlay, J. A. When anaerobes encounter oxygen: mechanisms of oxygen toxicity, tolerance and defence. Nature reviews microbiology 19, 774–785 (2021).

29 Van Acker, H. & Coenye, T. The Role of Reactive Oxygen Species in Antibiotic-Mediated Killing of Bacteria. Trends in microbiology 25, 456–466 (2017).

30 Krebs, N. F. Overview of zinc absorption and excretion in the human gastrointestinal tract. The Journal of nutrition 130, 1374S–1377S (2000).

31 Durand-Reville, T. F. et al. Rational design of a new antibiotic class for drug-resistant infections. Nature 597, 698–702 (2021).

32 Harwood, C. R. & Wipat, A. Sequencing and functional analysis of the genome of Bacillus subtilis strain 168. FEBS letters 389, 84–87 (1996).

33 Battye, T. G. G., Kontogiannis, L., Johnson, O., Powell, H. R. & Leslie, A. G. iMOSFLM: a new graphical interface for diffraction-image processing with MOSFLM. Acta Crystallographica Section D: Biological Crystallography 67, 271–281 (2011).

34 Evans, P. R. & Murshudov, G. N. How good are my data and what is the resolution? Acta Crystallographica Section D: Biological Crystallography 69, 1204–1214 (2013).

35 Winn, M. D. et al. Overview of the CCP4 suite and current developments. Acta Crystallographica Section D: Biological Crystallography 67, 235–242 (2011).

36 Vagin, A. A. & Lebedev, MoRDa, an automatic molecular replacement pipeline. Acta Crystallographica Section A: Foundations and Advances 71, S19 (2015).

37 Terwilliger, T. C. et al. Iterative model building, structure refinement and density modification with the PHENIX AutoBuild wizard. Acta Crystallographica Section D: Biological Crystallography 64, 61–69 (2008).

38 Liebschner, D. et al. Macromolecular structure determination using X-rays, neutrons and electrons: recent developments in Phenix. Acta Crystallographica Section D: Structural Biology 75, 861–877 (2019).

39 Emsley, P. & Cowtan, K. Coot: model-building tools for molecular graphics. Acta crystallographica section D: biological crystallography 60, 2126–2132 (2004).

40 Murshudov, G. N. et al. REFMAC5 for the refinement of macromolecular crystal structures. Acta Crystallographica Section D: Biological Crystallography 67, 355–367 (2011).

41 McGuffin, L. J., Bryson, K. & Jones, D. T. The PSIPRED protein structure prediction server. Bioinformatics 16, 404–405 (2000).

42 Rueden, C. T. et al. ImageJ2: ImageJ for the next generation of scientific image data. BMC Bioinformatics 18, 529 (2017).

43 Li, C. et al. Bis-Cyclic Guanidines as a Novel Class of Compounds Potent against Clostridium difficile. ChemMedChem 13, 1414–1420 (2018).

44 Edwards, A. N., Tamayo, R. & McBride, S. M. A novel regulator controls Clostridium difficile sporulation, motility and toxin production. Mol Microbiol 100, 954–971 (2016).

45 Edwards, A. N., Krall, E. G. & McBride, S. M. Strain-Dependent RstA Regulation of Clostridioides difficile Toxin Production and Sporulation. J Bacteriol 202, e00586–19 (2020).

46 Edwards, A. N. & McBride, S. M. Determination of the in vitro Sporulation Frequency of Clostridium difficile. Bio Protoc 7, e2125 (2017).

47 Ma, C. et al. Discovery of SARS-CoV-2 Papain-like Protease Inhibitors through a Combination of High-Throughput Screening and a FlipGFP-Based Reporter Assay. ACS Central Science 7, 1245–1260 (2021).

48 Marty, M. T. et al. Bayesian deconvolution of mass and ion mobility spectra: from binary interactions to polydisperse ensembles. Anal Chem 87, 4370–4376 (2015).

