## Extended Data for "A unique class of Zn^2+^-binding PBPs underlies cephalosporin resistance and sporogenesis of *Clostridioides difficile*"

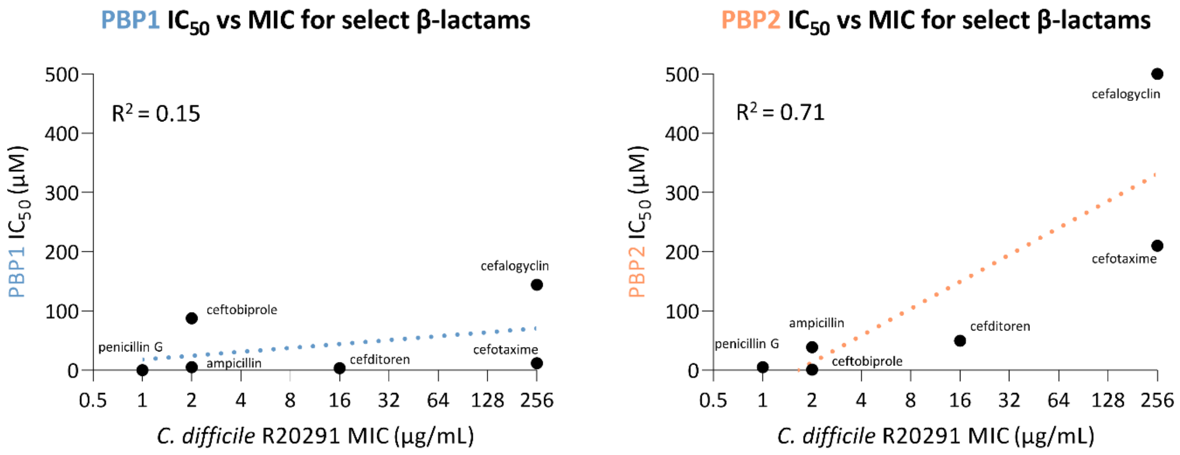


**Extended Data Fig. 1 | Relationship between $IC_{50}$ and antibacterial potency for select β-lactams against *C. difficile*.** Log correlation between $IC_{50}$ and MIC against *C. difficile* R20291 with residual plot shown as dashed line. β-Lactams include penicillin G, ampicillin, cefalogyclin, cefotaxime, cefditoren, and ceftobiprole; cefalexin was excluded as an outlier because $IC_{50}$ could not be determined.

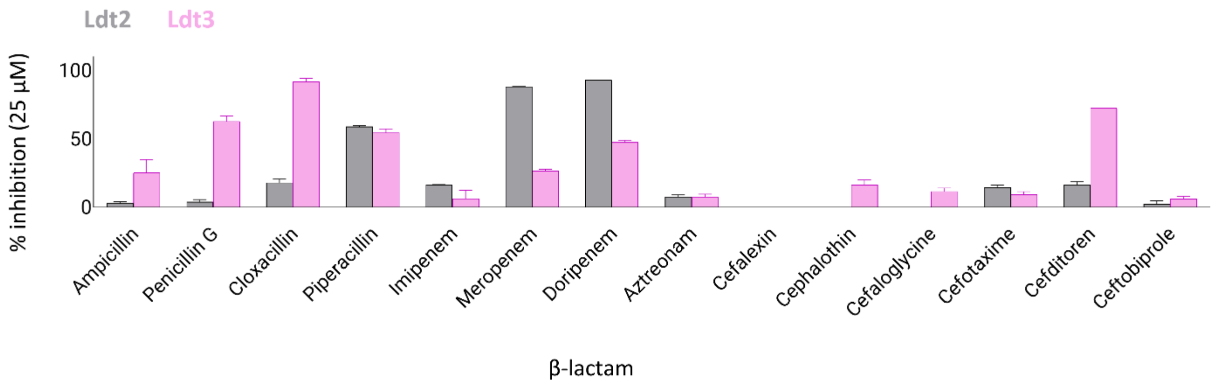


**Extended Data Fig. 2 | Inhibition of nitrocefin binding to Ldt2/Ldt3 by different β-lactam antibiotics at 25 µM.** β-lactams were weaker inhibitors of Ldt2 and Ldt3 compared to the tested PBPs. Meropenem, doripenem, and piperacillin were the best inhibitors of Ldt2, while cloxacillin and cefditoren were the best inhibitors of Ldt3.

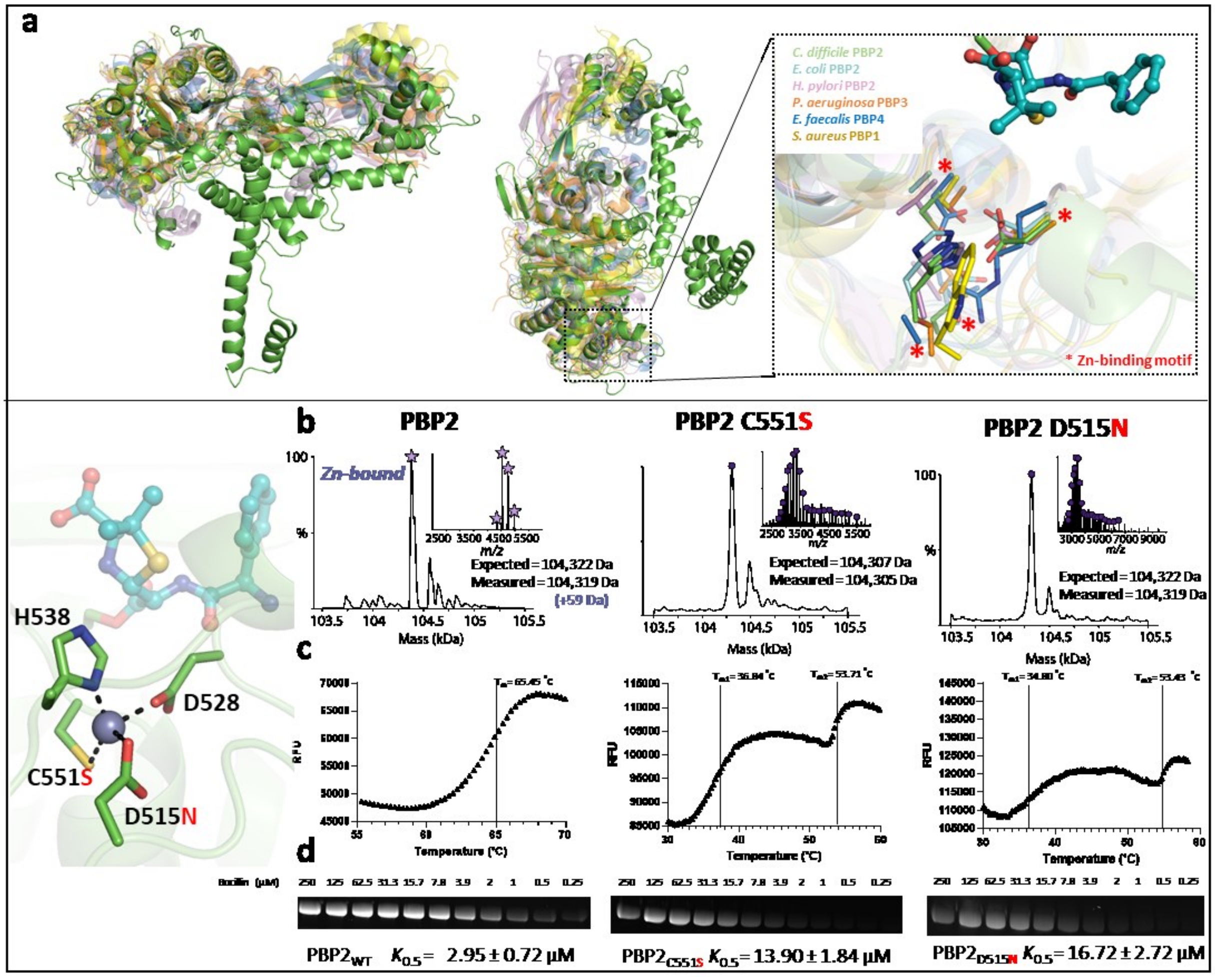


**Extended Data Fig. 3| A unique $Zn^{2+}$ -binding motif influences ligand binding and protein stability in PBP2. (a)** Structural superimposition of PBP2 with related homologues from select bacteria (left). The residues forming the $Zn^{2+}$ -binding motif from PBP2 are divergent amongst PBPs (right). **(b)** Native mass spectrometry, **(c)** bocillin titration, and **(d)** melting temperature assay for PBP2, PBP2 C551S, and PBP2 C551N. Note, the raw data shown is one representative of the triplicates performed.

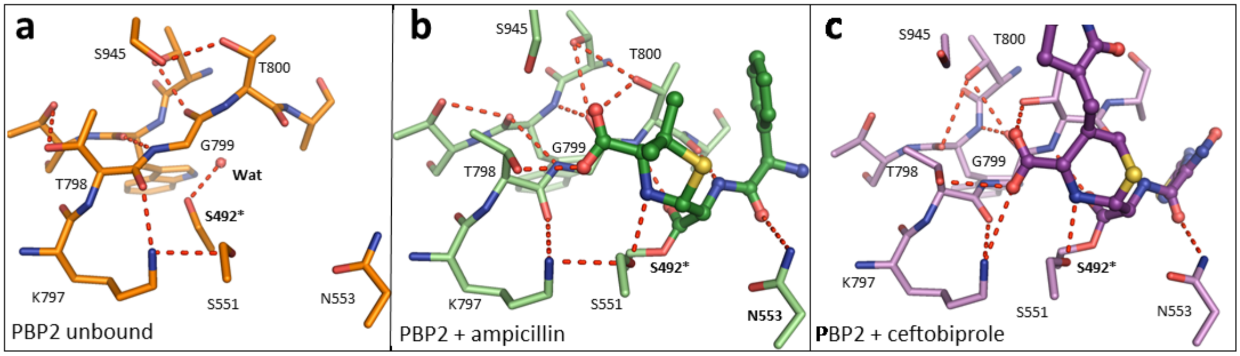


**Extended Data Fig. 4| Residues involved in ligand binding demonstrate structural rearrangement upon binding.** The catalytic core of PBP2 **(a)** unbound, **(b)** with ampicillin and **(c)** with ceftobiprole. Ligand binding induces conformational changes involving key residues, including the KTG motif, Asn553, and the catalytic Ser492*. These movements are accompanied by distinct hydrogen bonding patterns.

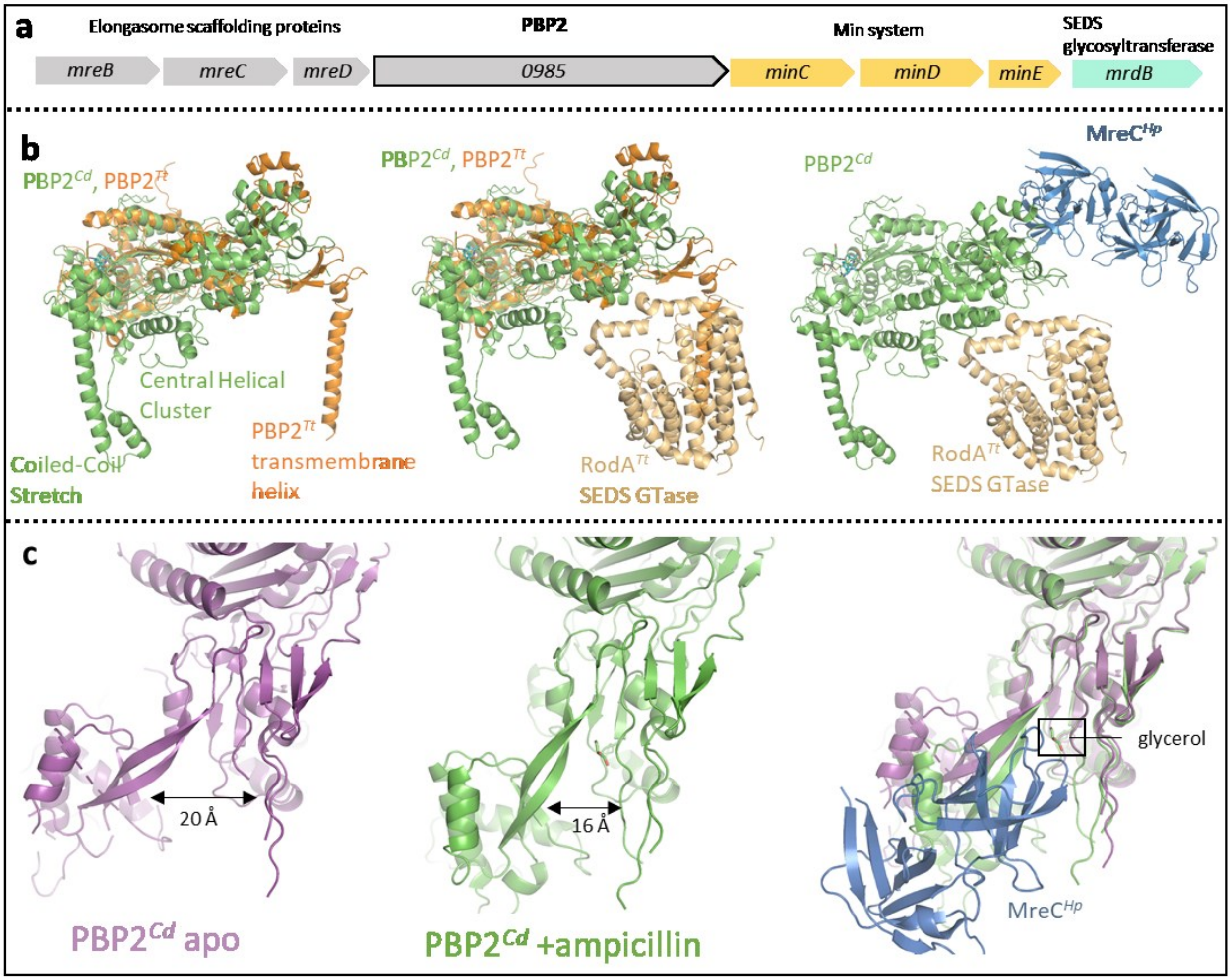


**Extended Data Fig. 5| Projected assembly for the peptidoglycan synthase of the elongasome.** **(a)** The gene encoding PBP2 is in a locus with several genes that are essential for cell elongation and division. PBP2 is the gene product of CDR20291_0985, the terminal gene of the mreBCD operon (*mreB*, *mreC*, *mreD*; grey). CDR20291_0985 is also adjacent to the genes encoding the Min system (*minC*, *minD*, *minE*; yellow), which facilitate the placement of the septal Z-ring. Because PBP2 is a monofunctional class B transpeptidase, it must rely on *mrdB* (teal), a transmembrane SEDS glycosyltransferase four genes downstream, to polymerize the sugar units of peptidoglycan. **(b)** Sideview of *C. difficile* PBP2 superimposed with *T. thermophilus* PBP2 (left, PDB 6PL5), *T. thermophilus* PBP2 + RodA (center, PDB 6PL5), and *T. thermophilus* RodA + H. pylori MreC (right, PDB 5LP5) **(c)** The NTD of apo (purple) and ampicillin-bound (green) PBP2 are splayed at different angles, resulting in a larger exposed crevasse for the apo-form. This region is putatively involved in the binding of MreC (right). In the ampicillin complex, there is electron density corresponding to a glycerol molecule where a portion of MreC (modelled from *H.pylori* MreC PDB 5LP5) normally binds.

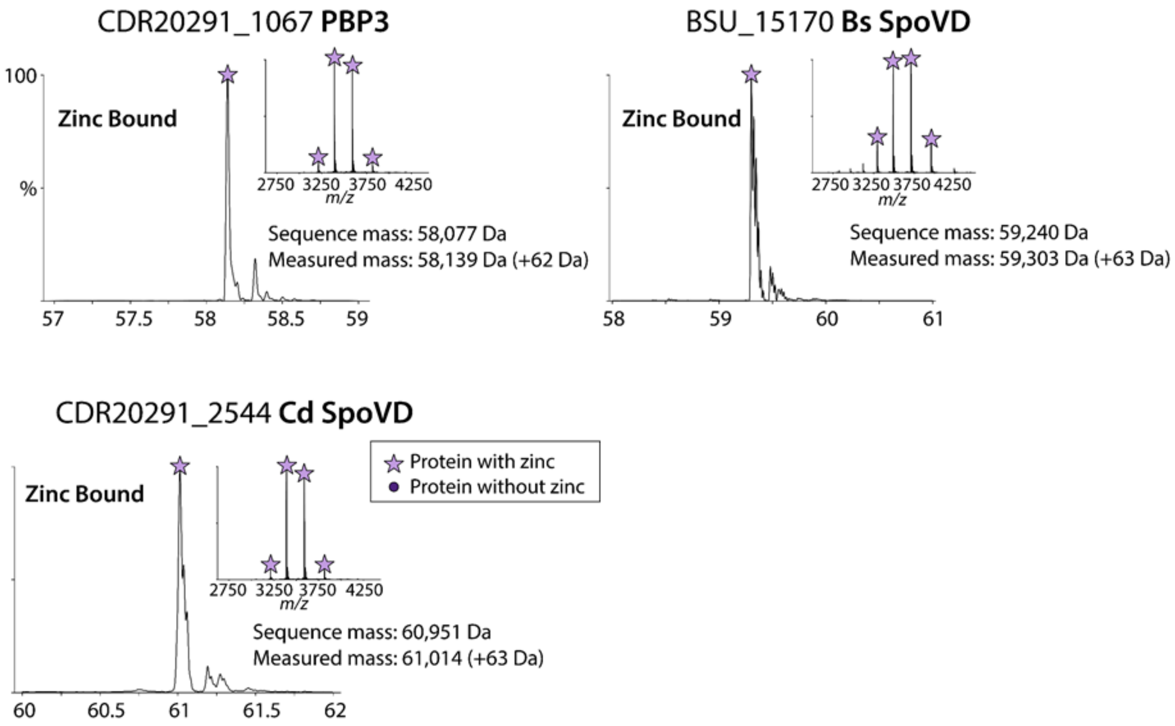


**Extended Data Fig. 6 |** Native mass spectrometry shows *C. difficile* SpoVD, PBP3, and *B. subtilis* SpoVD contain a mass peak corresponding to a $Zn^{2+}$-ion.

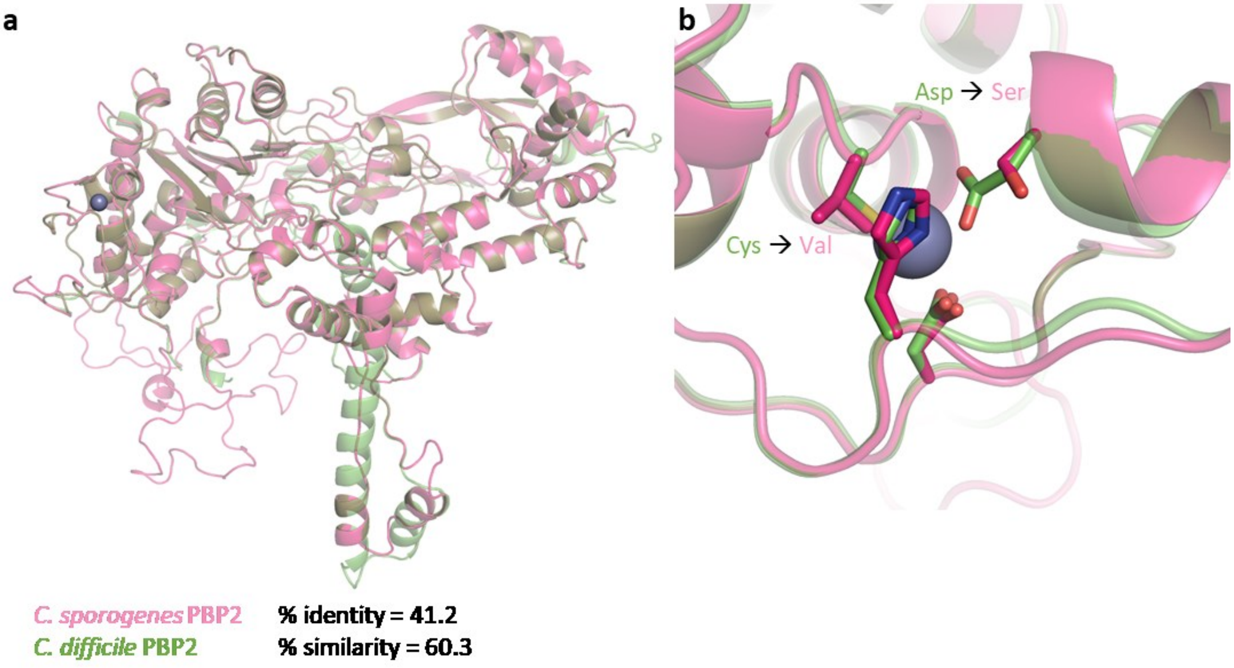


**Extended Data Fig. 7| *C. sporogenes* PBP2 bears high similarity to *C. difficile* PBP2 but lacks a $Zn^{2+}$ -binding motif. (a)** Structural superimposition of *C. difficile* PBP2 with homology model of *C. sporogenes* PBP2. *C. Sporogenes* is remarkably similar in identity, but lacks a coiled-coil stretch, and **(b)** residues capable of coordinating a $Zn^{2+}$ ion. *C. sporogenes* PBP2 does not appear to bind $Zn^{2+}$ due to the two substitutions in the motif.

| PDB ID | **7RCW** | **7RCX** | **7RCY** | **7RCZ** | **7RD0** |
|---|---|---|---|---|---|
| Gene (*R20291*) | *0985* | *0985* | *0985* | *2544* | *1067* |
| Protein | PBP2 | PBP2 | PBP2 | SpoVD | PBP3 |
| Inhibitor | Ampicillin | Apo | Ceftobiprole | Ampicillin | Apo |
| **Data Collection** | | | | | |
| Space Group | P 1 21 1 | P 1 21 1 | P 1 21 1 | P 21 21 21 | C 2 2 21 |
| Stoichiometry | 1mer | 1mer | 1mer | Homo 2mer | 1mer |
| Cell Dimensions | | | | | |
| a, b, c (Å) | 62.91, 200.19, 70.68 | 65.30, 122.46, 70.73 | 64.70, 196.39, 70.97 | 76.64, 95.62, 176.77 | 85.02 114.22, 154.63 |
| α, β, γ (°) | 90.00, 100.11, 90.00 | 90.00, 103.40, 90.00 | 90.00, 105.66, 90.00 | 90.00, 90.00, 90.00 | 90.00, 90.00, 90.00 |
| Resolution (Å) | 49.34 – 3.00 | 48.81 - 2.85 | 53.84 - 3.00 | 49.53 - 2.20 | 45.94 - 2.40 |
| | (3.16 – 3.00) | (3.00 - 2.85) | (3.11 - 3.00) | (2.25 - 2.20) | (2.49 - 2.40) |
| $R_{merge}$ | 0.103 (0.580) | 0.067 (0.406) | 0.067 (0.425) | 0.095 (0.461) | 0.078 (0.916) |
| <I>/σ<I> | 9.2 (2.4) | 10.6 (2.1) | 9.3 (2.0) | 7.3 (2.5) | 8.3 (2.0) |
| Completeness (%) | 90.2 (86.1) | 92.2 (86.0) | 83.9 (81.6) | 94.6 (96.8) | 99.9 (99.9) |
| CC1/2 (%) | 98.9 (77.4) | 99.4 (75.3) | 98.1 (67.5) | 98.5 (85.9) | 99.2 (74.3) |
| Redundancy | 3.6 (3.6) | 3.1 (2.9) | 2.6 (2.6) | 3.5 (3.4) | 5.7 (5.8) |
| **Refinement** | | | | | |
| Resolution (Å) | 49.34 – 3.00 | 39.78 - 2.85 | 53.84 – 3.00 | 42.05 - 2.20 | 42.51 - 2.40 |
| | (3.11 - 3.00) | (2.95 - 2.85) | (3.16 - 3.00) | (2.28 - 2.20) | (2.49 - 2.40) |
| No. reflections/free | 30828 / 1505 | 23206 / 1135 | 28434 / 1483 | 62337 / 3188 | 29753 / 1528 |
| $R_{work}/R_{free}$ | 0.208 / 0.257 | 0.220 / 0.275 | 0.215 / 0.269 | 0.193 / 0.228 | 0.194 / 0.236 |
| Clashscore | 7.81 | 9.14 | 10.78 | 4.28 | 3.07 |
| No. Atoms | | | | | |
| Overall | 6500 | 6377 | 6527 | 8389 | 3907 |
| Protein | 6430 | 6343 | 6479 | 7929 | 3823 |
| Ligand/Ion | 42 | 22 | 37 | 141 | 25 |
| Water | 28 | 12 | 11 | 319 | 59 |
| B-Factors (Å$^2$) | | | | | |
| Overall | 69.55 | 74.52 | 85.20 | 44.39 | 63.47 |
| Protein | 69.72 | 74.52 | 85.28 | 44.09 | 63.53 |
| Ligand/Ion | 59.87 | 91.57 | 87.85 | 76.14 | 84.72 |
| Solvent | 43.65 | 40.05 | 42.62 | 37.79 | 50.24 |
| RMS Deviations | | | | | |
| Bond Lengths (Å) | 0.014 | 0.014 | 0.014 | 0.014 | 0.014 |
| Bond Angles (°) | 1.93 | 1.80 | 1.83 | 1.87 | 1.93 |
| Ramachandran Favored (%) | 89.34 | 91.89 | 90.88 | 96.91 | 94.56 |
| Ramachandran Allowed (%) | 10.66 | 7.99 | 9.00 | 2.89 | 4.60 |
| Ramachandran Outliers (%) | 0.00 | 0.12 | 0.12 | 0.20 | 0.84 |
| Rotameric Outliers (%) | 3.58 | 4.06 | 5.26 | 2.43 | 2.08 |

**Extended Data Table 1 | Table of crystallization statistics**. Values in parentheses correspond to highest-resolution shell. All datasets were collected from a single crystal.

**Extended Data Table 2 | Bacteria with PBPs containing Zn-binding motifs.**

**Extended_Data_Table2.xlsx**

**Extended Data Table 3 | PBP proteins for phylogenetic analysis. PBP proteins containing a $Zn^{2+}$-binding motif are shaded red, those without are shaded blue**

Extended_Data_Table3.xlsx
